# Testing potential mechanisms of conspecific sperm precedence in *Drosophila pseudoobscura*

**DOI:** 10.1101/2021.02.09.430534

**Authors:** Brooke Peckenpaugh, Dean M. Castillo, Leonie C. Moyle

## Abstract

*Drosophila pseudoobscura* females that co-occur with sister species *D. persimilis* show elevated fertilization by conspecific sperm when they mate with both a heterospecific and a conspecific male. This phenomenon, known as conspecific sperm precedence (CSP), has evolved as a mechanism to avoid maladaptive hybridization with *D. persimilis*. In this study, we assessed pericopulatory (during mating) and postcopulatory (after mating) traits in crosses with sympatric or allopatric *D. pseudoobscura* females and conspecific or heterospecific males to evaluate potential mechanisms of CSP in this system. We observed shorter copulation duration in crosses with sympatric females, but found no difference in quantity of sperm transferred or female reproductive tract toxicity between sympatry and allopatry. Our data also support the hypothesis that parasperm, a short, sterile sperm morph, can protect fertile eusperm from the *D. pseudoobscura* female reproductive tract, though it is unclear how this might affect patterns of sperm use in sympatry vs. allopatry. Overall, these results suggest that copulation duration could potentially contribute to the elevated CSP observed in sympatry.

## Introduction

Among sexually reproducing organisms, the formation of new species ultimately requires the evolution of reproductive barriers. The resulting reproductive isolation is often associated with barriers that act before mating. However, costly hybridization can also be prevented through mechanisms that act after mating, including conspecific sperm precedence (CSP). CSP occurs when a female preferentially uses conspecific sperm to fertilize her eggs, after having mated with both a heterospecific and conspecific male. CSP can be a strong reproductive barrier among plant and animal species (Howard 1999, and references therein), including in *Drosophila* (Price 1997, Chang 2004, Castillo and Moyle 2019). However, the specific mechanisms that underlie this female postcopulatory preference for conspecific sperm are currently poorly understood (Price et al. 2000), except in the most intensely studied model systems (Manier et al. 2013a, b).

CSP could be due to mechanisms that act during mating, or after mating has occurred but prior to fertilization. In the first instance, two pericopulatory (during mating) mechanisms that can influence fertilization patterns are variation in copulation duration and in the amount of sperm and/or other ejaculate components that are transferred. Of these, copulation duration is known to vary among species and among crosses (Lizé et al. 2011), and longer copulation duration is associated with a greater delay in female remating (Gilchrist and Partridge 2000), which can increase the fertilization success of the associated male by reducing the opportunity for subsequent sperm competitors. This effect is likely the consequence of increased transfer of seminal fluid proteins (Wigby et al. 2009), as longer copulation duration has been associated with greater transfer of sex peptide and ovulin—seminal fluid proteins known to stimulate ovulation, elevate egg production, and decrease female receptivity to remating (Chapman et al. 2003, Rubinstein and Wolfner 2013). Copulation duration might also be important for appropriately stimulating female postmating fertilization responses; in particular, the mechanical stimulation of copulation during conspecific matings has been shown to induce females to eject previously stored sperm (Snook and Hosken 2004), although the specific importance of copulation duration for this effect is yet to be examined. Finally, both sexes affect copulation duration to some extent (Spieth 1974, Hirai et al. 1999, Lefranc and Bundgaard 2000, Mazzi et al. 2009), indicating that it could respond to selection acting on females, males, or both sexes. For example, *Drosophila* males can make sophisticated decisions about seminal fluid protein allocation according to female condition and mating status (Sirot 2019, Sirot et al. 2011), so this could be a mechanism by which males preserve reproductive resources when mating with unpreferred (for example, heterospecific) females.

Similar to copulation duration, sperm transfer during copulation can also influence fertilization patterns. In addition to directly determining the amount of sperm available for storage by females, sperm transfer is also known to be mediated by male reproductive allocation decisions. For example, *Drosophila* males can modulate sperm transfer according to social context (i.e., the presence of rivals) (Sirot 2019, Price et al. 2012, Garbaczewska et al. 2013), or the perceived quality of females (Lüpold et al. 2010). Interestingly, although copulation duration has been shown to positively correlate with the number of sperm transferred in some insect species (e.g., melon fly, Tsubaki and Sokei 1988; scorpionfly, Engqvist and Sauer 2003), there is little evidence for this in *Drosophila* (Gilchrist and Partridge 2000, Price et al. 2012), suggesting that these two traits might have somewhat independent effects on fertilization patterns, with copulation duration primarily affecting seminal fluid rather than sperm transfer *per se*. Regardless, given their properties, these pericopulatory mechanisms could contribute to CSP if heterospecific matings are shorter, and/or if males transfer fewer sperm to heterospecific females, resulting in proportionally more conspecific sperm available in storage.

Mechanisms of CSP could also act after copulation and sperm transfer, via primarily female-mediated postmating traits. For example, *Drosophila* females can bias sperm use through sperm storage patterns (Manier et al. 2013c), or through the timing of sperm ejection after remating (Manier et al. 2013a). The postcopulatory mechanism of particular interest in our study is female reproductive tract toxicity. Holman and Snook (2008) demonstrated that the female reproductive tract (FRT) can be toxic to sperm after conspecific *D. pseudoobscura* matings. This toxicity might enhance female fitness if it reduces the frequency of low quality or unsuitable sperm in the FRT, thereby acting as a filter on sperm quality (Holman and Snook 2006). While FRT toxicity has previously only been assessed within species (Holman and Snook 2008), this phenomenon could contribute to CSP if female reproductive tracts also preferentially kill heterospecific sperm over conspecific sperm. Under this scenario, multiply mated females could reduce fertilization from low quality heterospecific males while retaining sufficient sperm from higher quality conspecific males. Finally, in addition to FRT toxicity, shifts in fertilization patterns can also be influenced by some additional male-mediated postcopulatory processes. For example, species in the *Drosophila obscura* group have heteromorphic sperm—males produce both a long, fertile morph (eusperm) and a short, sterile morph (parasperm); sperm heteromorphism has been proposed to influence patterns of sperm competition within species (Snook 1998, Wedell and Cook 1999, Oppliger et al. 2003). Nonetheless, since the specific function of parasperm is not yet well understood, and they do not contribute directly to fertilization (Snook and Karr 1998), it is difficult to predict *a priori* whether the type of sperm transferred could potentially also play a role in CSP.

While reproductive barriers such as CSP often evolve as a byproduct of sexual dynamics within species, selection can also actively drive their formation. For example, when geographically overlapping species are insufficiently diverged to prevent hybridization, selection can act to increase reproductive barriers that reduce the formation of lower fitness hybrids—a process called “reinforcement” (Dobzhansky 1940). When reinforcement occurs, reproductive isolation is expected to be elevated specifically in sympatry (in direct response to selection to prevent hybridization), but not allopatry (where there is no contact, and consequently no selection pressure). Therefore, one can demonstrate evidence for reinforcement by comparing relevant phenotypes in sympatry versus allopatry. Although the generality of reinforcement in driving speciation in nature is not yet known, several compelling examples have previously documented this pattern (e.g., Coyne and Orr 1989, Noor 1995, Saetre et al. 1997, Hopkins and Rausher 2012). Moreover, while reinforcement is commonly associated with selection on premating isolating barriers, it can also act after mating (e.g., Matute 2010). In particular, Castillo and Moyle (2019) demonstrated that CSP has responded to reinforcing selection in *Drosophila pseudoobscura* where it co-occurs with *D. persimilis*. Specifically, sympatric *D. pseudoobscura* females, upon mating with both *D. persimilis* and *D. pseudoobscura* males, use more conspecific sperm than allopatric females do. The mechanism(s) of CSP in this system is currently unknown.

In this study, we sought to evaluate (a) two potential pericopulatory mechanisms of CSP (copulation duration, and sperm transfer), (b) a potential postcopulatory mechanism of CSP (female reproductive tract toxicity), and (c) a potential parasperm function in *D. pseudoobscura* and *D. persimilis*. Since CSP is observed to be elevated in sympatric *D. pseudoobscura* females compared to allopatric females (Castillo and Moyle 2019), if these mechanisms play a role in reinforced CSP we expected to observe shorter copulation between *D. persimilis* males and sympatric *D. pseudoobscura* females, fewer heterospecific sperm transferred to sympatric females, and/or higher rates of heterospecific sperm death specifically in sympatric female reproductive tracts. Finally, we expected ejaculates with higher parasperm proportions to be associated with increased eusperm viability, as previously demonstrated (Holman and Snook 2008). Although the relationship with CSP is unclear a priori, we can also evaluate whether parasperm patterns vary between allopatry and sympatry. To assess these expectations, we evaluated both heterospecific and conspecific crosses, using *D. pseudoobscura* females from sympatric and allopatric populations, to compare these traits across populations and cross types.

## Methods

### Fly stocks

All stocks were reared on standard media prepared by the Bloomington Drosophila Stock Center and were kept at room temperature (~22°C). We used a set of isofemale lines collected from four natural populations in the summers of 2013 and 2014, and previously examined for CSP in Castillo and Moyle (2019). Allopatric *D. pseudoobscura* were collected at Zion National Park, UT (provided by N. Phadnis) and Lamoille Canyon, NV (collected by D. Castillo). Sympatric *D. pseudoobscura* and *D. persimilis* were collected at two sites: Mt. St. Helena, CA (*D. pseudoobscura* collected by A. Hish/M. Noor and D. Castillo, and *D. persimilis* collected by D. Castillo); and, near Meadow Vista and Forest Hill, CA (called here ‘Sierra’; *D. pseudoobscura* and *D. persimilis* collected by D. Castillo). National Drosophila Species Stock Center identification numbers for these lines can be found in Table S1. For both sympatric populations, both species were present in field collections and can therefore be considered co-occurring.

### Experimental matings to assess peri- and post-copulatory variation

Of the 16 *D. pseudoobscura* lines analyzed in Castillo and Moyle (2019), up to eight isofemale lines—two from each of Zion and Lamoille (allopatric), and Mt. St. Helena and Sierra (sympatric)—were used as experimental female lines. To compare patterns of trait variation in sympatric versus allopatric female lines, we performed crosses between isofemale *D. pseudoobscura* lines from each population and males of two types: heterospecific males (up to two lines from each *D. persimilis* sympatric location); or conspecific males (from the same isofemale line as the experimental female). In each case, we isolated females and males and allowed them to age for seven days before setting up a cross. Male virgins were set up in mating vials 24 hours before each mating to allow them to acclimate. A female was then added, mating observed, and latency to copulation and copulation duration recorded (in minutes). (Throughout, we detected no differences in latency to copulation between cross types, so those data are not further considered here.) Following mating, females were either sacrificed immediately (control treatments) or retained for a set period of time (FRT exposure treatments; either 30 minutes or 2 hours, depending upon the specific experiment) prior to being sacrificed. After sacrificing, each female’s reproductive tract (bursa, spermathecae, and seminal receptacle) removed and dissected. In each sample, postcopulatory sperm viability and sperm identity (eusperm versus parasperm) data was quantified as detailed below. Control and FRT exposure treatment crosses for each cross type were completed in parallel (on the same day whenever possible) and in a randomized order.

### Heterospecific and conspecific sperm viability assay (two hours of FRT exposure)

We assessed both heterospecific and conspecific sperm traits after two hours of FRT exposure in each of four *D. pseudoobscura* female lines, one from each of the four geographical sites—Zion and Lamoille (allopatric), and Mt. St. Helena and Sierra (sympatric) (Table 1). For heterospecific matings, we mated each female line to a sympatric *D. persimilis* line from Mt. St. Helena (the same line as used in 30 min FRT assays). For conspecific matings, we mated each female to a male from her own line. We completed four technical replicates for each cross type (n = 32, for each of heterospecific and conspecific crosses). In all cases, we set up each cross with one female and two males of the target identity, as the presence of two males helped to facilitate heterospecific matings.

**Table 1.**
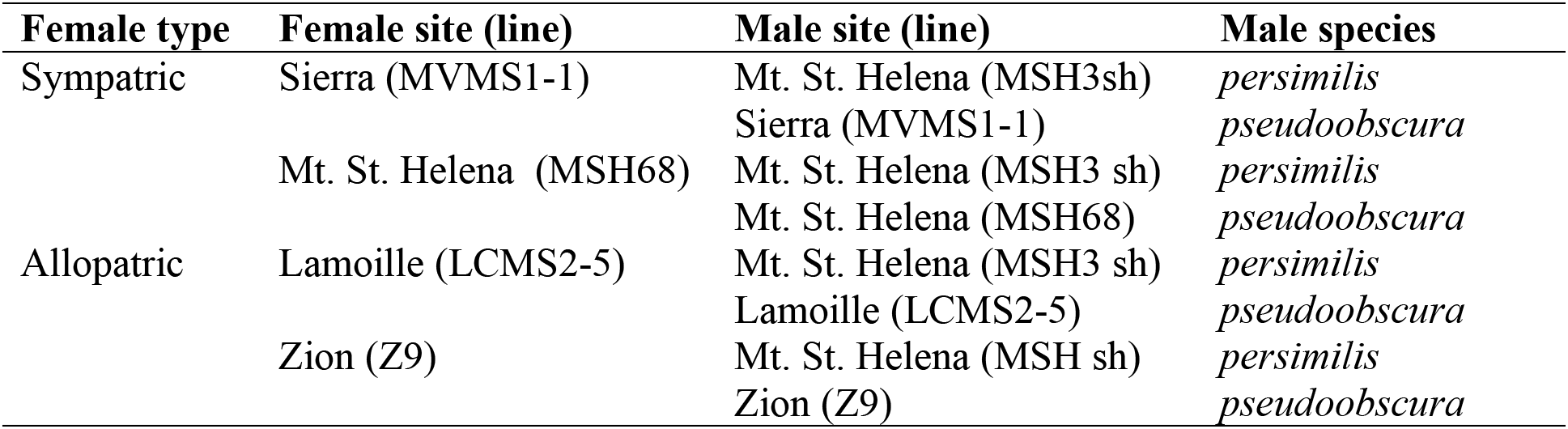
All cross combinations and line names used in two hour exposure matings.

### Heterospecific sperm viability assay (30 minutes of FRT exposure)

In addition to 2 hour FRT exposure for both con- and heterospecific males, we also assessed just heterospecific sperm viability after 30 minutes of FRT exposure in order to evaluate evidence for viability effects across a larger number of different female lines. For these assays, we used eight *D. pseudoobscura* female lines, two from each of the four geographical sites— Zion and Lamoille (allopatric), and Mt. St. Helena and Sierra (sympatric) (Table S1). Females from each of these lines were crossed to each of two *D. persimilis* male lines, one from Mt. St.

Helena and one from Sierra (both sympatric sites). We completed three technical replicates for each cross type (n = 96 total crosses).

### Scoring sperm viability, eusperm/parasperm proportion, and amount of sperm transferred

We followed the sperm viability assessment protocol from Holman and Snook (2008). For each FRT exposure treatment, the ejaculate was left in the female for the designated exposure time (either 30 minutes or two hours) before dissecting the reproductive tract into 20 μL of room temperature Grace’s insect medium, and assessing sperm traits. For each control cross, the female reproductive tract was dissected into 20 μL of room temperature Grace’s insect medium immediately after mating; the sample was then left covered and in the dark for either 30 minutes or two hours, prior to assessing sperm traits. This ensured that ‘control’ samples were the same age as treatment samples, and differed only in whether they experienced prolonged exposure to the FRT. For all samples, immediately prior to preparation and imaging, clumps of sperm were dispersed with a pipette and the sample was divided in half (10 μL each). One half was prepared for viability staining and quantifying sperm transfer, and the other half was used for eusperm/parasperm classification.

To classify eusperm and parasperm proportions, we stained one half of each sample (10 μL) with SpermBlue (Microptic) and took digital images under the EVOS FL microscope for blind counting later. SpermBlue enables clear visualization of entire sperm including their tails, allowing for reliable classification of sperm morphs based on tail length; these features cannot be clearly discerned from the viability stain, which only stains the sperm head.

To classify live versus dead sperm in the remaining half (10 μL) of each sample, we prepared samples by adding 1 μL of dilute LIVE/DEAD sperm viability stain (Molecular Probes L-7011; 1 μL SYBR-14, 2 μL propidium iodide, 45.5 μL Grace’s insect medium), and left the stained sample for five minutes before transferring it to a hemocytometer for counting. Images of the live/dead stained sperm were taken under an EVOS FL digital inverted fluorescence microscope and labeled with numbers for blind counting later. The quantity of sperm transferred was counted from a 0.1 μL sample (a corner square of the hemocytometer, selected at random); all sperm transfer numbers reported are therefore means per 0.1 μL sample. Sperm viability was determined by counting the number of dead sperm in the same sample and subtracting that from the total. Individual matings that did not transfer a detectable amount of sperm were excluded from this dataset and analysis; specifically, out of 188 total matings, 20 attempts were excluded due to lack of sperm (n = 11 sympatric, n = 9 allopatric).

Finally, to further assess estimates of sperm viability in the absence of FRT exposure, we also quantified sperm viability from a subset of cross types, by dissecting sperm either directly from the male testes (see supplementary methods, Figure S1) or immediately after mating (without sitting in Grace’s insect medium for two hours (see supplementary methods, Figure S2).

### Statistical analyses

All analyses were completed in R version 4.0.2.

#### i) Differences in copulation duration

We assessed evidence for reduced copulation duration in sympatry, and whether this differed between conspecific and heterospecific matings. Because the premating setup is identical for all treatments, we used data from all crosses for this analysis (two hour exposure, 30 min exposure, and controls). We used a linear mixed model, with female type (sympatric vs. allopatric) and male species (*D. pseudoobscura* vs. *D. persimilis*) as fixed effects, the interaction between female type and male species as a fixed interaction effect, and female line as a random effect, using the lmer function (Bates et al. 2015). We included the interaction term to identify potential differences in the female response based on the species of male with which she mated. Significance was determined using the Satterthwaite approximation, implemented in the lmerTest package (Kuznetsova et al. 2017).

#### ii) Differences in quantity of sperm transferred

We evaluated evidence for reduced sperm transfer in sympatry, whether this effect is specifically observed with heterospecific males, and whether copulation duration predicts the quantity of sperm transferred. We only used control data for this analysis (where sperm is dissected from females immediately after mating), to avoid any effect on sperm counts that might otherwise result from prolonged exposure to the female reproductive tract (e.g., the storage of sperm, or incorporation into the mating plug). We used a linear mixed model, with female type (sympatric vs. allopatric), male species (*D. pseudoobscura* vs. *D. persimilis*), and copulation duration (a covariate) included as fixed effects, and female line included as a random effect, using the lmer function (Bates et al. 2015). Significance was determined using the Satterthwaite approximation, implemented in the lmerTest package (Kuznetsova et al. 2017).

#### iii) Differences in patterns of female reproductive tract toxicity

We evaluated evidence for differences in eusperm viability following prolonged FRT exposure (two hours) compared to immediate dissection and equivalent aging in Grace’s insect medium, including whether this difference depended on female or male identity. Live versus dead eusperm is a binomial response, so we used a generalized linear mixed model where female type (sympatric vs. allopatric), male species (*D. pseudoobscura* vs. *D. persimilis*), and treatment group (two hour exposure to the FRT or to Grace’s insect medium) were included as fixed effects, and female line was included as a random effect. We implemented this model using the glmer function, which assesses significance using a Wald test (Bates et al. 2015; see Table 2).

**Table 2.** Generalized linear mixed model results assessing patterns in eusperm viability. Female line was also included as a random effect.

| Model Component | Estimate | Std. Error | z | p |
| --- | --- | --- | --- | --- |
| Intercept | 0.825 | 0.110 | 7.499 | < 0.0001 |
| Female type (sympatric) | 0.052 | 0.126 | 0.412 | 0.680 |
| Male species (*D. persimilis*) | -0.009 | 0.104 | -0.087 | 0.931 |
| Treatment group (two hour FRT exposure) | 0.341 | 0.105 | 3.24 | **0.001** |

#### iv) Effect of parasperm proportion on eusperm viability

To evaluate whether eusperm viability after exposure to the FRT is influenced by the proportion of parasperm within the same ejaculate sample, we used a generalized linear mixed model with live and dead eusperm counts coded as a binomial response. We used data from two hour treatment and control crosses for this analysis (Table 1). The parasperm proportion data are taken from SpermBlue assays (see above), which are counted from the same sample as the eusperm viability data. We included female type (sympatric vs. allopatric), male species (*D. pseudoobscura* vs. *D. persimilis*), treatment group (two hour exposure to the FRT or to Grace’s insect medium), and parasperm proportion as fixed main effects, the interaction between treatment group and parasperm proportion as a fixed interaction effect, and female line as a random effect. We included the interaction term to identify potential differences in the effect of parasperm proportion on the response between treatment and control groups. We implemented this model using the glmer function, which assesses significance using a Wald test (Bates et al. 2015).

## Results

### (a) Shorter copulation duration could contribute to elevated CSP in sympatry

We first sought to assess whether pericopulatory traits—specifically copulation duration and/or amount of sperm transferred—could contribute to the observed decrease in heterospecific sperm use in sympatry. We found that female type (sympatric or allopatric) significantly predicts copulation duration (LMM, *β* = 3.22, *P* = 0.002). In particular, copulation is shorter with sympatric females compared to allopatric females (Figure 1A). In contrast, there is no difference in copulation duration between heterospecific and conspecific males (LMM, *β* = 0.899, *P* = 0.925), and there is no significant interaction effect between female type and male species (LMM, *β* = −2.12, *P* = 0.118). Overall, these results suggest that copulation duration is primarily moderated by females, regardless of male type. The general pattern of shorter copulation duration in sympatry indicates that this trait could contribute to reduced heterospecific sperm use in these females.

**Figure 1.**
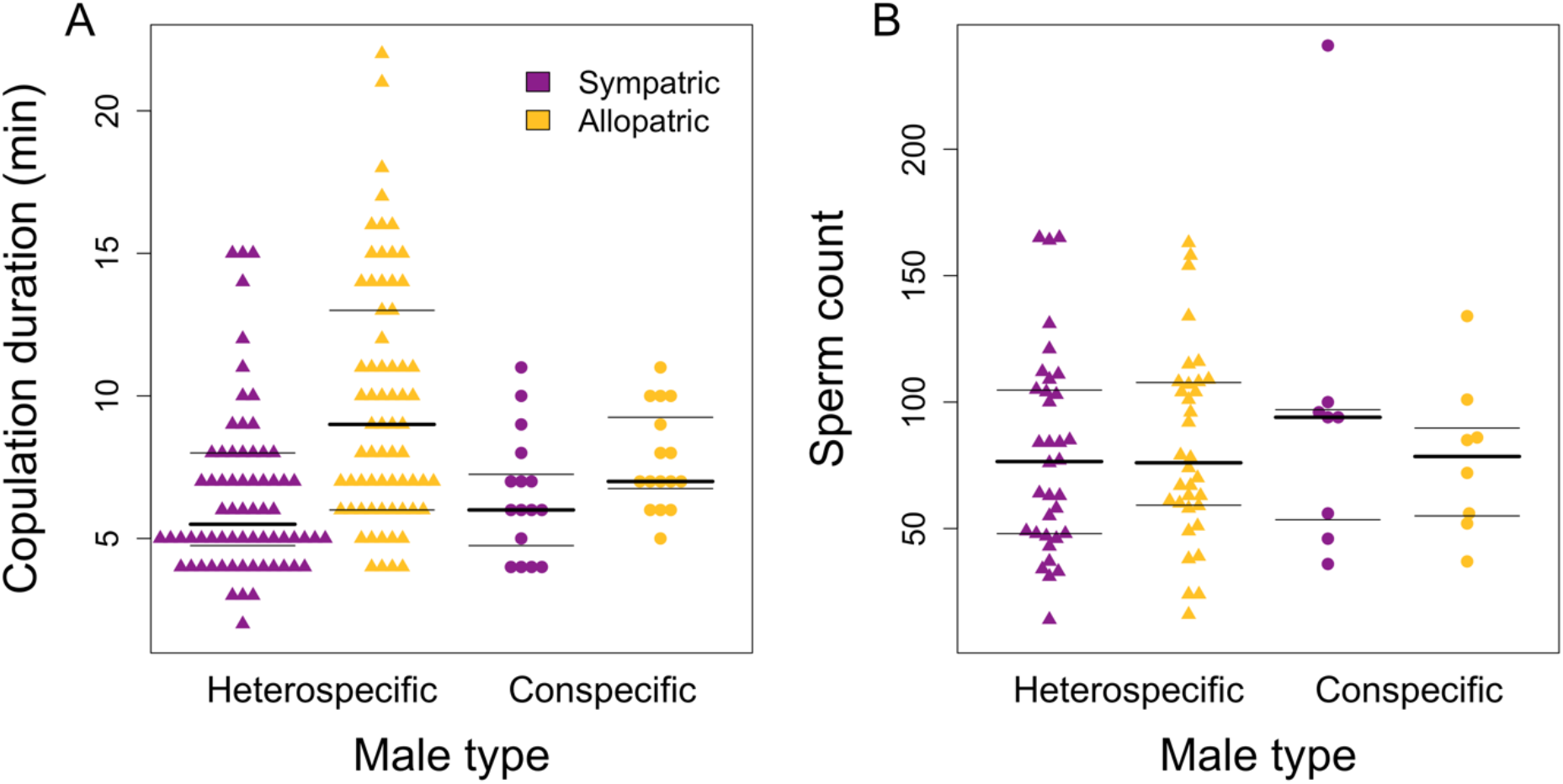
(A) Copulation duration differs between sympatric and allopatric females (LMM, *β* = −2.86, *P* = 0.005). (B) Quantity of sperm transferred does not differ depending on female type (LMM, *β* = 3.40, *P* = 0.808) or male species identity (*β* = −10.37, *P* = 0.354). The thicker plot line indicates the median, the thinner lines indicate quartiles for each group.

Unlike copulation duration, neither female type nor male species predict the quantity of sperm transferred (Figure 1B; LMM, *β* = 3.40, *P* = 0.808 and *β* = −10.37, *P* = 0.354, respectively). Furthermore, copulation duration does not predict the number of sperm transferred (LMM, *β* = 1.19, *P* = 0.081). This result indicates that reduced heterospecific sperm use in sympatric females is unlikely to be explained by a specific reduction in heterospecific sperm transfer. We only used control data to estimate the quantity of sperm transferred, since sperm quantity differed significantly between treatment and control groups after two hours of FRT exposure (t-test, *t* = 2.53, *P* = 0.014)—that is, it was systemically lower in the FRT treatment— presumably due to sperm storage, ejection, or recruitment to the mating plug in that time.

### (b) Female reproductive tract toxicity does not explain elevated CSP

We next assessed whether postcopulatory trait variation—specifically elevated FRT toxicity—could contribute to the observed decrease in heterospecific sperm use in sympatry. Reduced eusperm viability after two hours of FRT exposure, specifically in sympatric heterospecific matings, would support this hypothesis. However, we instead found significantly greater eusperm viability in the treatment (two hours of FRT exposure) compared to the control (GLMM, *β* = 0.341, *P* = 0.001). This viability difference could be due to elevated sperm death in the controls, since our data show that a significant proportion of sperm die after two hours of exposure to Grace’s insect medium (Figure S1, Figure S2). Additionally, it could be due to the preferential loss of dead sperm after two hours in the FRT treatment (i.e., ejected or retained in the mating plug), or to both effects. In either case, we found no difference in eusperm viability associated with female type (sympatric vs. allopatric—*β* = 0.052, *P* = 0.680). Finally, eusperm viability did not differ significantly according to male species (*β* = −0.009, *P* = 0.931) or among individual female lines (Wald test, χ^2^ = 6.3, *P* = 0.097; see Figure S3). Thus, patterns of eusperm viability are not consistent with a role for female reproductive tract toxicity in the reduced use of *D. persimilis* sperm by sympatric *D. pseudoobscura* females (Figure 2).

**Figure 2.**
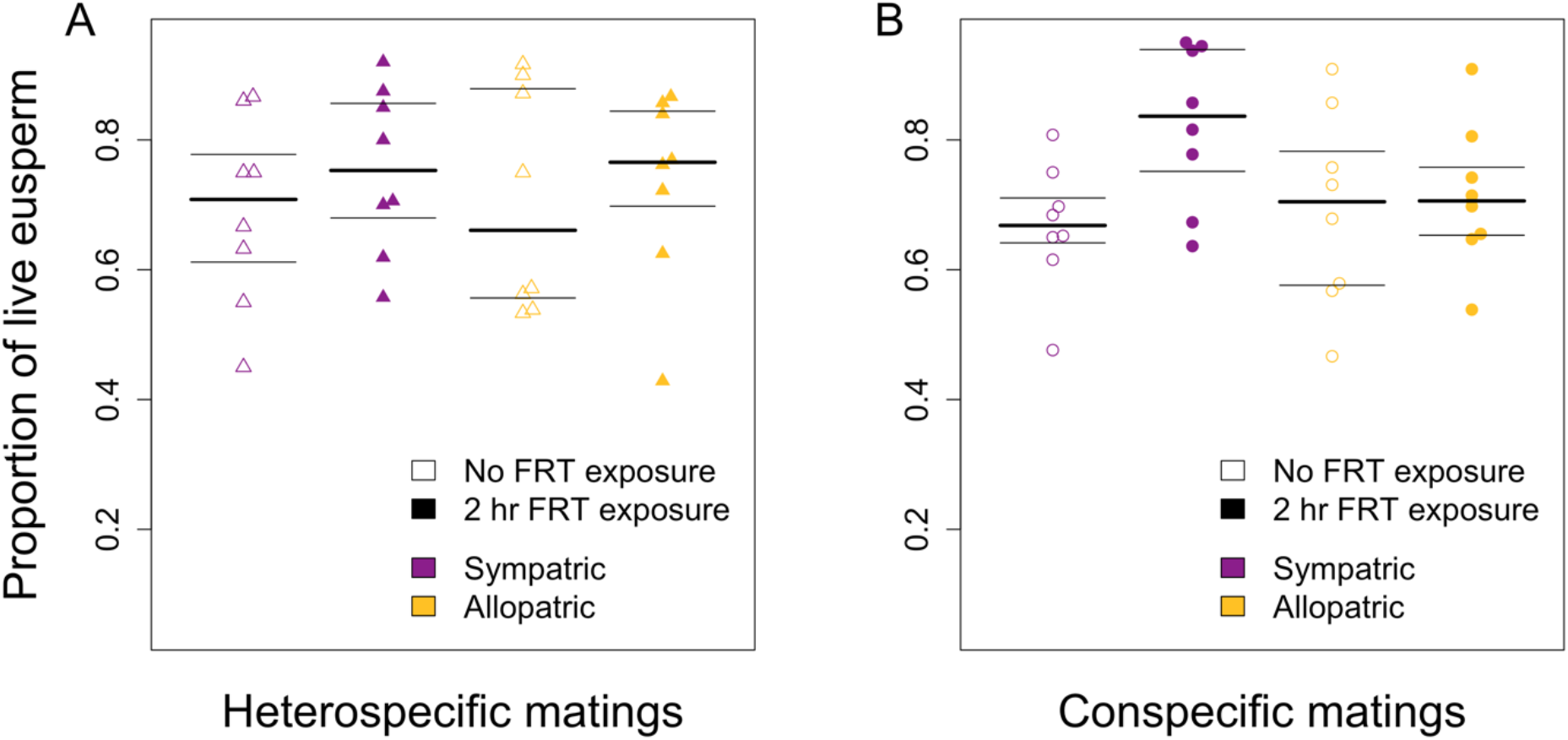
Eusperm viability differs between 0 hour FRT exposure controls and 2 hour FRT exposure crosses (GLMM, *β* = 0.341, *P* = 0.001) in matings with (A) *D. persimilis* males and (B) *D. pseudoobscura* males. The thicker plot line indicates the median, the thinner lines indicate quartiles for each group.

### (c) Parasperm proportion predicts eusperm viability after FRT exposure

Finally, we evaluated whether higher parasperm proportion in an ejaculate is associated with increased eusperm viability after exposure to the FRT. We found a significant interaction effect between the proportion of parasperm and treatment group (GLMM, *β* = 1.50, *P* = 0.044), which reflects a positive relationship between parasperm proportion and eusperm viability only in the FRT treatment group (Figure 3). This is consistent with previous findings, suggesting that parasperm proportion predicts eusperm viability in the FRT. Parasperm proportion does not impact eusperm viability without FRT exposure (*β* = −0.128, *P* = 0.769; Figure 3) and there is no difference in the proportion of parasperm between treatment groups (*β* = −0.145, *P* = 0.611). Similarly, female type (*β* = 0.069, *P* = 0.631) and male species *(β* = −0.054, *P* = 0.623) do not significantly predict eusperm viability. This result suggests that parasperm could play a role in protecting eusperm in the FRT, that does not depend on specific genotypic effects or geographical context.

**Figure 3.**
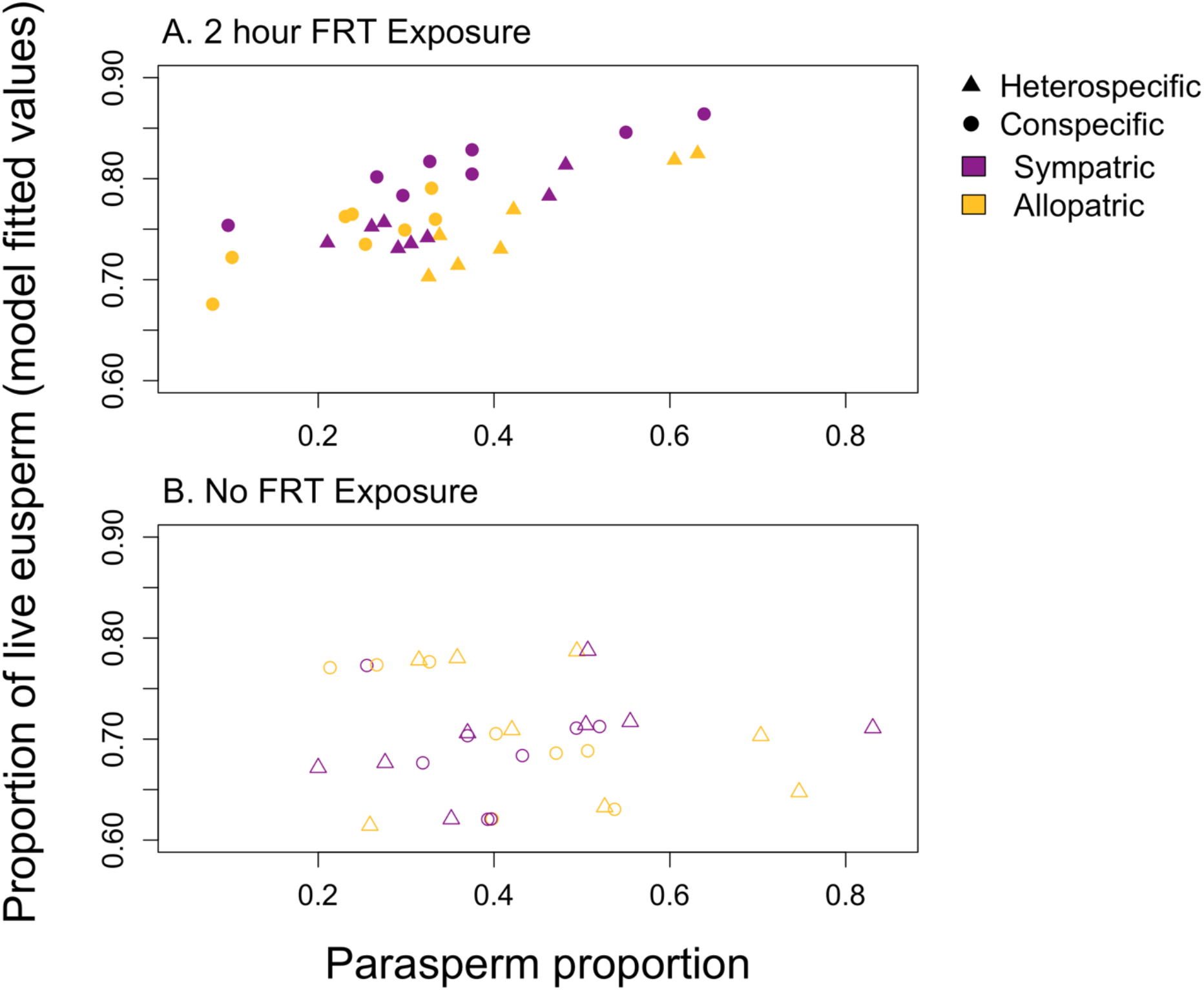
(A) Parasperm proportion predicts eusperm viability after two hours of FRT exposure (GLMM, *β* = 1.50, *P* = 0.044). (B) Parasperm proportion does not predict eusperm viability in control crosses, where sperm is dissected immediately after mating. Triangles represent heterospecific matings, circles represent conspecific matings.

## Discussion

Conspecific sperm precedence can be an important barrier to maladaptive hybridization in nature (Howard et al. 1998, Chang 2004, Price 1997), and is known to have experienced reinforcing selection where the sister species *D. pseudoobscura* and *D. persimilis* co-occur (Castillo and Moyle 2019). In this study, we assessed several potential peri- and post-copulatory mechanisms that could contribute to this reduced heterospecific sperm use observed specifically in sympatric *D. pseudoobsura* females. Of two pericopulatory mechanisms, we found that patterns of copulation duration, but not sperm transfer, are consistent with a possible role in elevated CSP in sympatry. Of postcopulatory mechanisms—specifically FRT toxicity—we found complex patterns of sperm viability after FRT exposure that suggest this mechanism cannot explain reinforced CSP. However, our data support previous findings that parasperm could play a role in protecting eusperm in the FRT. Our results exclude several peri- and postcopulatory mechanisms as targets of reinforcing selection in this case, but indicate that reproductive responses mediated by copulation duration, or other mechanisms not measured here, could play an important role in reducing maladaptive hybridization via CSP in this species pair.

### Pericopulatory mechanisms of reinforced CSP

We found that copulation duration is shorter between heterospecific males and sympatric females compared to matings with allopatric females, as might be predicted if copulation duration had responded to reinforcing selection. Our observation that copulation duration is not quantitatively associated with the amount of sperm transferred (as also observed in other species—Price et al. 2012, Gilchrist and Partridge 2000) indicates that the potential effect of shorter copulation on reduced paternity share in sympatry is not via direct effects on reduced sperm transfer. However, copulation duration could influence paternity share via other mechanisms. For instance, longer copulations facilitate the transfer of more seminal fluid proteins (Wigby et al. 2009), which can influence paternity share by increasing female refractoriness to remating (Laturney et al. 2018) or by increasing sperm competitive ability, sperm transport and storage, egg-laying behavior, and more (Chapman and Davies 2004). Previous analyses indicate that fractoriness to remating does not differ between sympatric and allopatric matings in our populations of *D. pseudoobscura* (Davis et al. 2016) and is therefore not likely to contribute to reinforcement of CSP here, but the other aforementioned effects of seminal fluid proteins remain to be tested. Furthermore, length of copulation might be important for the mechanical stimulation of female postmating fertilization responses, (Snook and Hosken 2004), with longer copulation duration producing more effective stimulatory responses. Overall, because pericopulatory mechanisms act early in reproduction to determine the ratio of conspecific to heterospecific sperm that ends up in storage, they need not require additional complex postcopulatory mechanisms, such as the sophisticated shifting of sperm use between sperm storage organs shown in *D. simulans* (Manier et al. 2013b). Thus, pericopulatory mechanisms could potentially evolve more easily. Regardless of mechanism, our results suggest that reduced copulation duration in sympatric *D. pseudoobscura* females could explain the reduced use of heterospecific sperm in sympatry.

Despite the fact that many previous studies of pericopulatory mechanisms show evidence for male mediation (Sirot et al. 2011, Price et al. 2012, Garbaczewska et al. 2013, Lüpold et al. 2010), our data suggest that this shift in copulation duration may be primarily mediated by a general response in sympatric females. In principle, a shift in copulation duration could be mediated by *D. persimilis* males in response to sympatric *D. pseudoobscura* females; for example this could act to reduce their investment in maladaptive heterospecific copulations, similar to how *D. melanogaster* males are able to adjust ejaculate size based on mate quality (Lüpold et al. 2010). Nonetheless, a primarily male-mediated response is not strongly supported by our observations. Not only is copulation duration not quantitatively associated with the amount of sperm transferred in our dataset, but we find that sympatric females have reduced copulation duration with males regardless of whether they are heterospecific or conspecific (Figure 1A). Theory suggests that females often bear the greater burden from low quality matings in nature (Trivers 1972), so terminating copulation early could be an adaptive mechanism for mitigating this evolutionary burden in sympatry. Overall, then, our results suggest that reduced copulation duration in sympatric females could contribute to the elevated

CSP observed in sympatry, although the specific mechanism by which this differentially influences paternity share in heterospecific males remains to be determined.

### Postcopulatory mechanisms of reinforced CSP

Postcopulatory mechanisms are also known to play a role in biased female sperm use (e.g. Snook and Hosken 2004, Manier et al. 2013a, b, c). In this study we specifically tested FRT toxicity as a potential new postcopulatory mechanism of CSP by evaluating whether more heterospecific sperm die in the FRT of sympatric female lines, compared to the FRT of allopatric female lines. Our results indicate that sperm death is not elevated in heterospecific, sympatric matings. In fact, we found that the proportion of dead sperm is, on average, reduced after exposure to the FRT. The greater proportion of viable sperm in the two hour FRT exposure treatment group could have a technical and/or a biological explanation. First, sperm die over time in Grace’s insect medium (Figure S1, Figure S2, Guo et al. 2021), suggesting that the FRT is a less toxic environment for sperm than the control medium. Furthermore, we see a reduction in the total number of sperm between control and treatment groups after two hours of FRT exposure. It is therefore also possible that dead sperm are being preferentially ejected or amassed into the mating plug over the duration of time spent in the FRT, leading to a higher proportion of viable sperm after two hours. Regardless, our results indicate that FRT toxicity against heterospecific sperm does not serve as a barrier to hybridization in this species, and further, has not responded to reinforcing selection in sympatry.

We also assessed sperm viability from conspecific matings in order to understand FRT toxicity in *D. pseudoobscura* females more generally. As with heterospecific crosses, we did not find consistent FRT toxicity against sperm. Data from across all female lines in our experiment indicate that there is considerable genetic variation for this female trait both among and within populations of *D. pseudoobscura* that could be available to respond to selection (Figure S4). Nonetheless, our results suggest that FRT toxicity is not a general reproductive response in *D. pseudoobscura* females.

While our analysis of parasperm does not explain patterns of CSP in this species, parasperm proportion data showed interesting patterns nonetheless. The function of parasperm— including its role in mediating sperm competition and use—is not currently well understood, though several hypotheses have been proposed for its evolution in the *obscura* group. For instance, parasperm have been suggested to displace a rival male’s eusperm from the sperm storage organs, or depress female remating behavior by filling her sperm storage organs (the “cheap filler” hypothesis) (Snook 1998). Furthermore, Holman and Snook (2008) proposed that parasperm act to buffer their co-transferred eusperm from the toxic female reproductive tract, thereby increasing sperm competitiveness and storage of direct genetic relatives. Our results support this hypothesis, since ejaculates with higher proportions of parasperm also show elevated viability of eusperm (Figure 3). This pattern cannot explain CSP, since it does not differ between sympatry and allopatry, but it does lend further support to this hypothesis for parasperm function.

In general, then, we did not find evidence for FRT toxicity as a postcopulatory mechanism for elevated CSP. This observation does not exclude the potential involvement of other post-copulatory mechanisms in reinforcing responses. For instance, the timing of sperm ejection can determine the composition of the ejaculate in the female reproductive tract (Snook and Hosken 2004, Manier et al. 2013b), and is known to vary between *Drosophila* species (Manier et al. 2013a). Furthermore, females of some *Drosophila* species can exert more sophisticated postcopulatory control of paternity through biased patterns of sperm storage and use (Manier et al. 2013c). Therefore mechanisms like sperm ejection time and sperm storage patterns could be relevant to explaining reinforced CSP in this system.

## Conclusion

Overall, of the peri- and post-copulatory mechanisms examined here, our data suggest that reduced copulation duration could contribute to elevated CSP between *D. pseudoobscura* and *D. persimilis* in sympatry. In contrast, our data suggest that neither patterns of heterospecific sperm transfer nor female reproductive tract toxicity are consistent with a role in reinforced CSP in sympatric *D. pseudoobscura*. Nonetheless, we found support for parasperm-mediated protection of eusperm while in the *pseudoobscura* FRT, regardless of the geographical origin of females or the species of males. Future work could identify the specific effect of copulation duration on patterns of heterospecific sperm use, as well as any interactions between copulation duration and other CSP mechanisms.

## Acknowledgements

We thank Nitin Phadnis, Alexander J. Hish, and Mohamed A. F. Noor for providing fly lines used in this study, and Donn Castillo for help with collecting strains. We thank Luke Holman and Rhonda R. Snook for advice on the sperm viability protocol. Research was supported by Indiana University Department of Biology funding to LCM and BP.

## Notes

**Conflict of interest statement** The authors have no conflict of interest to declare.

### Competing Interest Statement

The authors have declared no competing interest.

