## Supplementary Materials for "Testing potential mechanisms of conspecific sperm precedence in *Drosophila pseudoobscura*"

**Supplementary Methods**

In order to interpret patterns of sperm death in the reported data, including the point at which most sperm death is occurring, we also estimated sperm viability at two additional (earlier) stages: after dissection directly from the seminal vesicles within the testes, and immediately after mating, without any waiting period (30 minutes or two hours) following dissection. For each assay, *D. pseudoobscura* and *D. persimilis* males were isolated and aged for seven days, as were *D. pseudoobscura* females in the second assay.

*Sperm viability in the testes*

We dissected and quantified sperm from testes of mature, unmated males, from each of six male lines (every male line that had been used in crosses previously): four *D. pseudoobscura* lines (Zion and Lamoille (allopatric), Mt. St. Helena and Sierra (sympatric)), and two *D. persimilis* lines from Sierra and Mt. St. Helena. We completed three technical replicates for each line (n = 18 total dissections). We dissected the testes into 10 μL of Grace's insect medium, then dispersed clumps of sperm with a pipette. The sample was stained and prepared for counting as described above in "Scoring sperm viability".

*Sperm viability immediately after mating*

We evaluated patterns of viability and sperm morph abundance in both heterospecific and conspecific crosses immediately after copulation using two female lines, one from allopatry (Lamoille) and one from sympatry (Mt St Helena) (the same isofemale lines used in assays above). For conspecific matings, we mated each female to a male from her own line. For heterospecific matings, we mated each female line to a sympatric *D. persimilis* line from Mt. St. Helena (the same line used in assays reported in the main text). We completed three technical replicates for each cross type (n = 12 total crosses). We set up each cross as described in the main text, observed matings, and recorded mating latency and duration. However, rather than leaving the sample covered and kept in the dark for 30 minutes or two hours, as in previous controls, we stained and prepared the samples for counting immediately after dissecting.

**Supplementary Tables and Figures**

**Table S1.** Line numbers for the lines used in this study from the National Drosophila Species Stock Center at Cornell University.

| Species | Site | Line | SKU |
|---|---|---|---|
| *D. pseudoobscura* | Lamoille | LC5 | 14011-0121.280 |
| | | LCMS2-5 | 14011-0121.283 |
| | Mt. St. Helena | MSH1 | 14011-0121.286 |
| | | MSH68 | 14011-0121.288 |
| | Sierra | S4 | 14011-0121.274 |
| | | MVMS1-1 | 14011-0121.278 |
| | Zion | Z11 | 14011-0121.292 |
| | | Z9 | 14011-0121.296 |
| *D. persimilis* | Mt. St. Helena | MSH3 sh | 14011-0111.66 |
| | Sierra | FH2 sh | 14011-0111.65 |

**Table S2.** All cross combinations and line names used for 30 minute FRT exposure matings. All 30 minute FRT assessments use heterospecific (*D. persimilis*) male lines.

| Female type | Female site | Female line | Male site (line) |
|---|---|---|---|
| Sympatric | Sierra | MVMS1-1 | Mt. St. Helena (MSH3 sh) |
| | | | Sierra (FH2 sh) |
| | | S4 | Mt. St. Helena (MSH3 sh) |
| | | | Sierra (FH2 sh) |
| | Mt. St. Helena | MSH68 | Mt. St. Helena (MSH3 sh) |
| | | | Sierra (FH2 sh) |
| | | MSH1 | Mt. St. Helena (MSH3 sh) |
| | | | Sierra (FH2 sh) |
| Allopatric | Lamoille | LCMS2-5 | Mt. St. Helena (MSH3 sh) |
| | | | Sierra (FH2 sh) |
| | | LC5 | Mt. St. Helena (MSH3 sh) |
| | | | Sierra (FH2 sh) |
| | Zion | Z9 | Mt. St. Helena (MSH3 sh) |
| | | | Sierra (FH2 sh) |
| | | Z11 | Mt. St. Helena (MSH3 sh) |
| | | | Sierra (FH2 sh) |

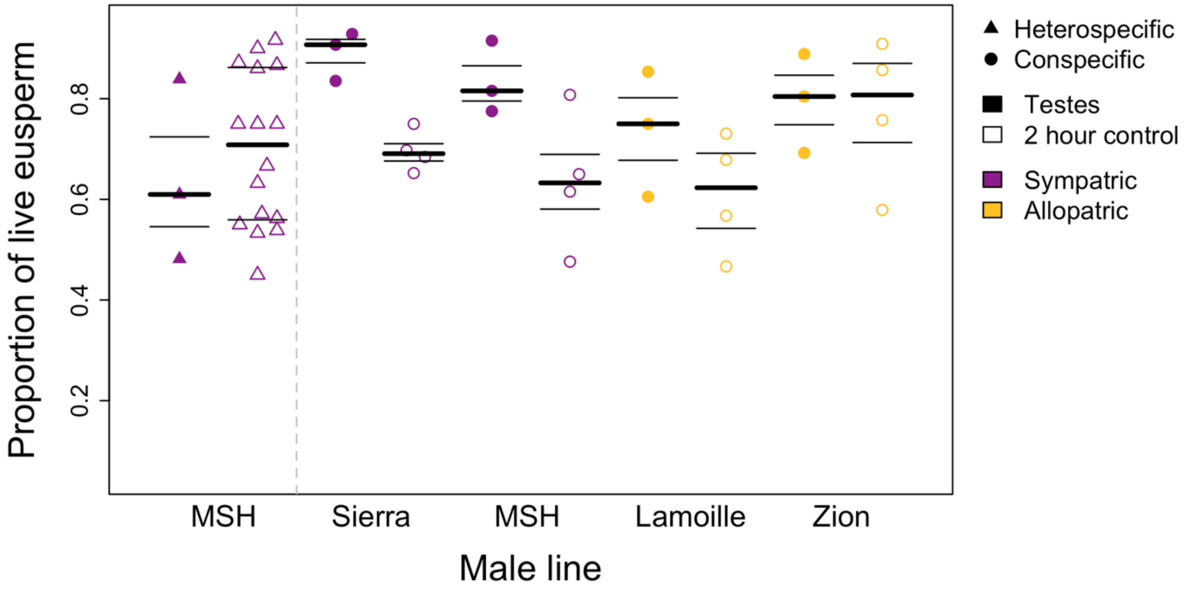


**Figure S1.** Eusperm viability is significantly reduced in the two hours they are sitting in media compared to immediately after dissection from the testes (GLMM, $\beta = 0.458$, $P < 0.001$). Closed symbols represent samples taken from testes and immediately prepared for counting, open symbols represent controls as seen in main text and Figure S1 (2 hr in Grace's insect medium). The thicker plot line indicates the median, the thinner lines indicate quartiles for each group.

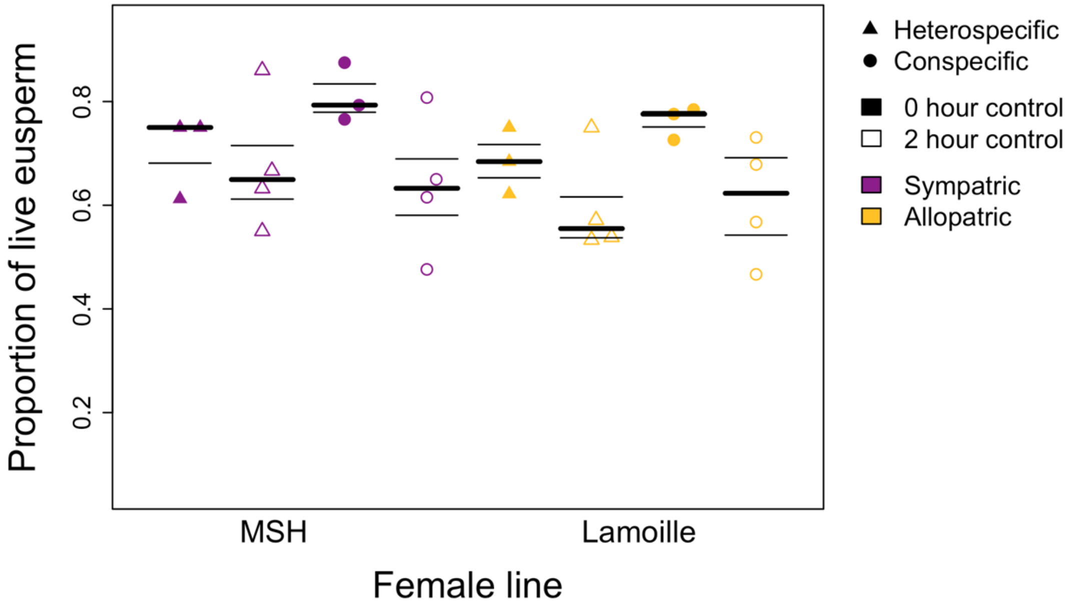


**Figure S2.** Eusperm viability is significantly reduced in the two hours they are sitting in media compared to immediately after mating (GLMM, $\beta = 0.466$, $P < 0.001$). Closed symbols represent samples counted immediately after dissection from the mated female, open symbols represent controls as seen in main text and Figure S1 (2 hr in Grace's insect medium). The thicker plot line indicates the median, the thinner lines indicate quartiles for each group.

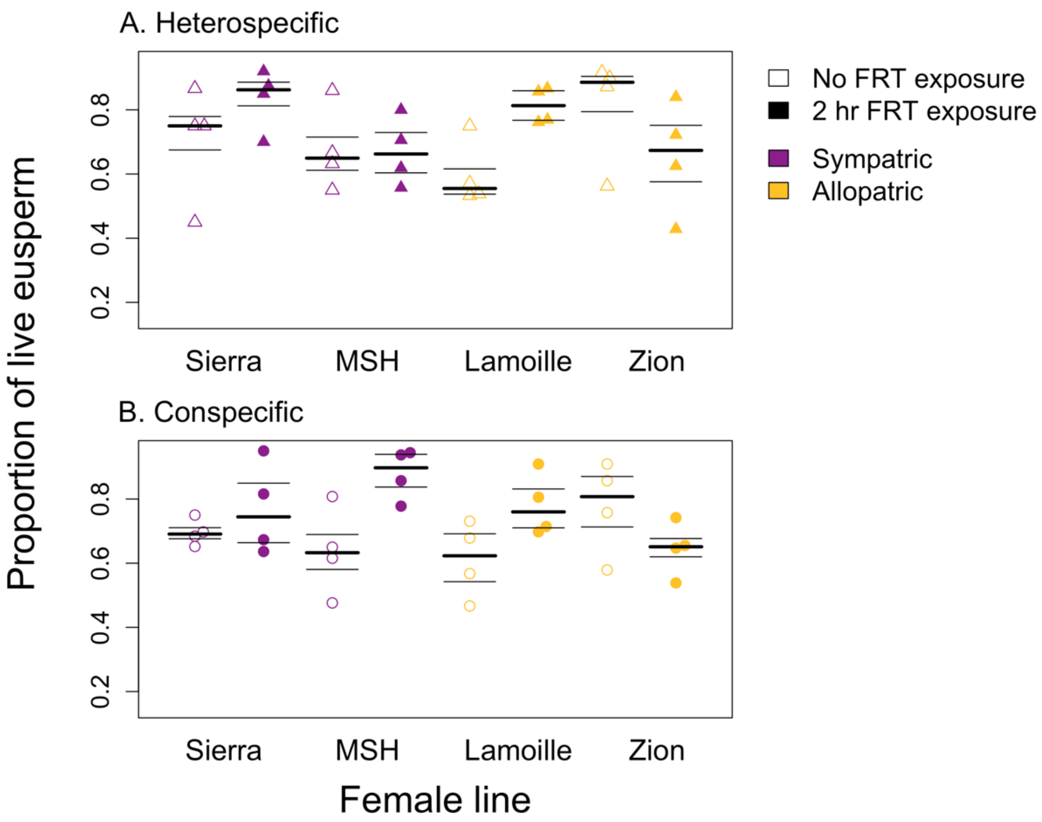


**Figure S3.** Patterns of eusperm viability did not vary significantly among individual female lines (Wald test, $\chi^2 = 6.3$, $P = 0.097$). Open symbols represent controls (2 hours in Grace's insect medium), closed symbols represent FRT exposed treatment (2 hours in FRT). The thicker plot line indicates the median, the thinner lines indicate quartiles for each group.

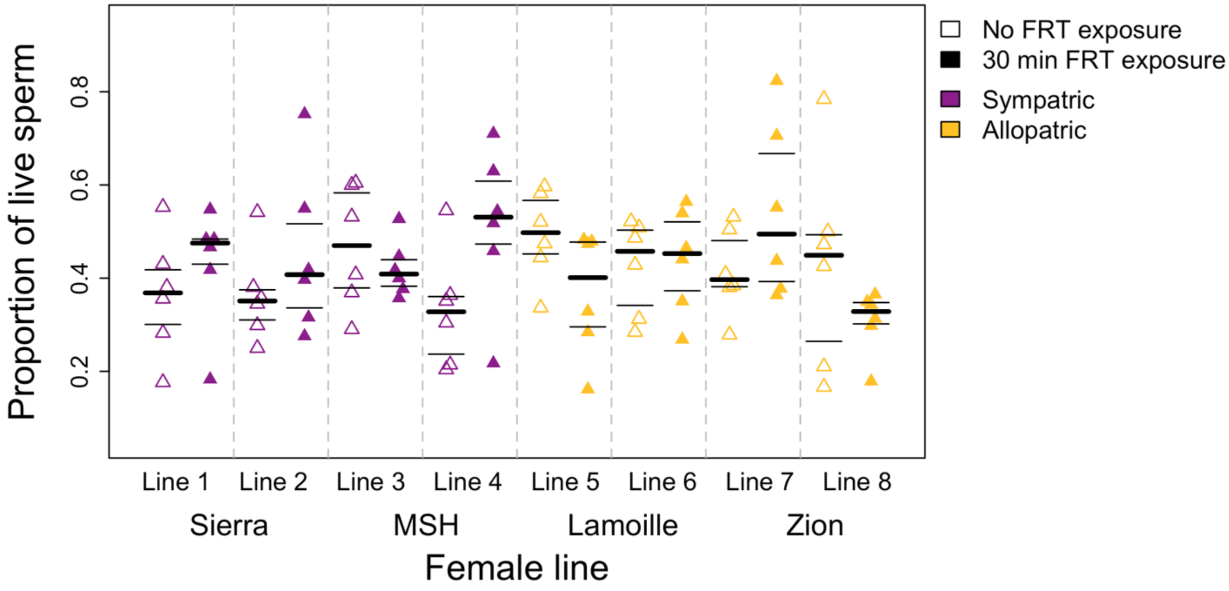


**Figure S4.** Total sperm (eusperm and parasperm) viability patterns vary significantly across female lines in heterospecific matings (Wald test, $\chi^2 = 12.4$, $P = 0.006$). Open symbols represent controls (30 min in Grace's insect medium), closed symbols represent FRT exposed treatment (30 min in FRT). The thicker plot line indicates the median, the thinner lines indicate quartiles for each group.

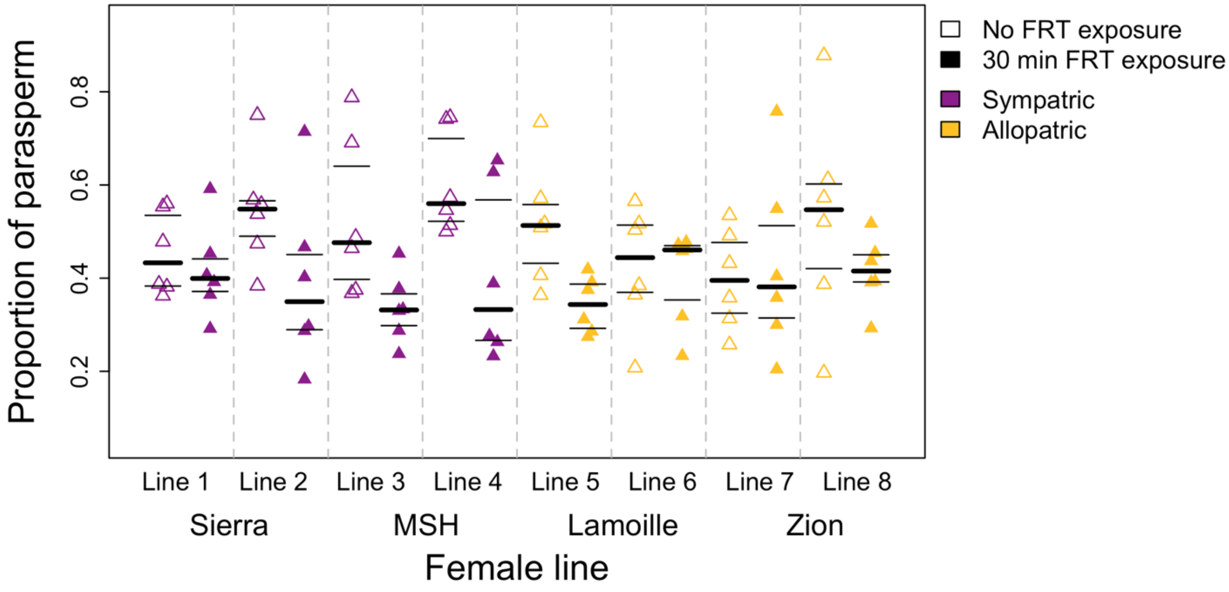


**Figure S5.** Parasperm proportion varies significantly across female lines in heterospecific matings (Wald test, $\chi^2 = 33.9$, $P < 0.001$). Crosses also have significantly less parasperm after two hours of FRT exposure (GLMM, $\beta = 0.443$, $P < 0.001$). Open symbols represent controls (30 min in Grace's insect medium), closed symbols represent FRT exposed treatment (30 min in FRT). The thicker plot line indicates the median, the thinner lines indicate quartiles for each group.

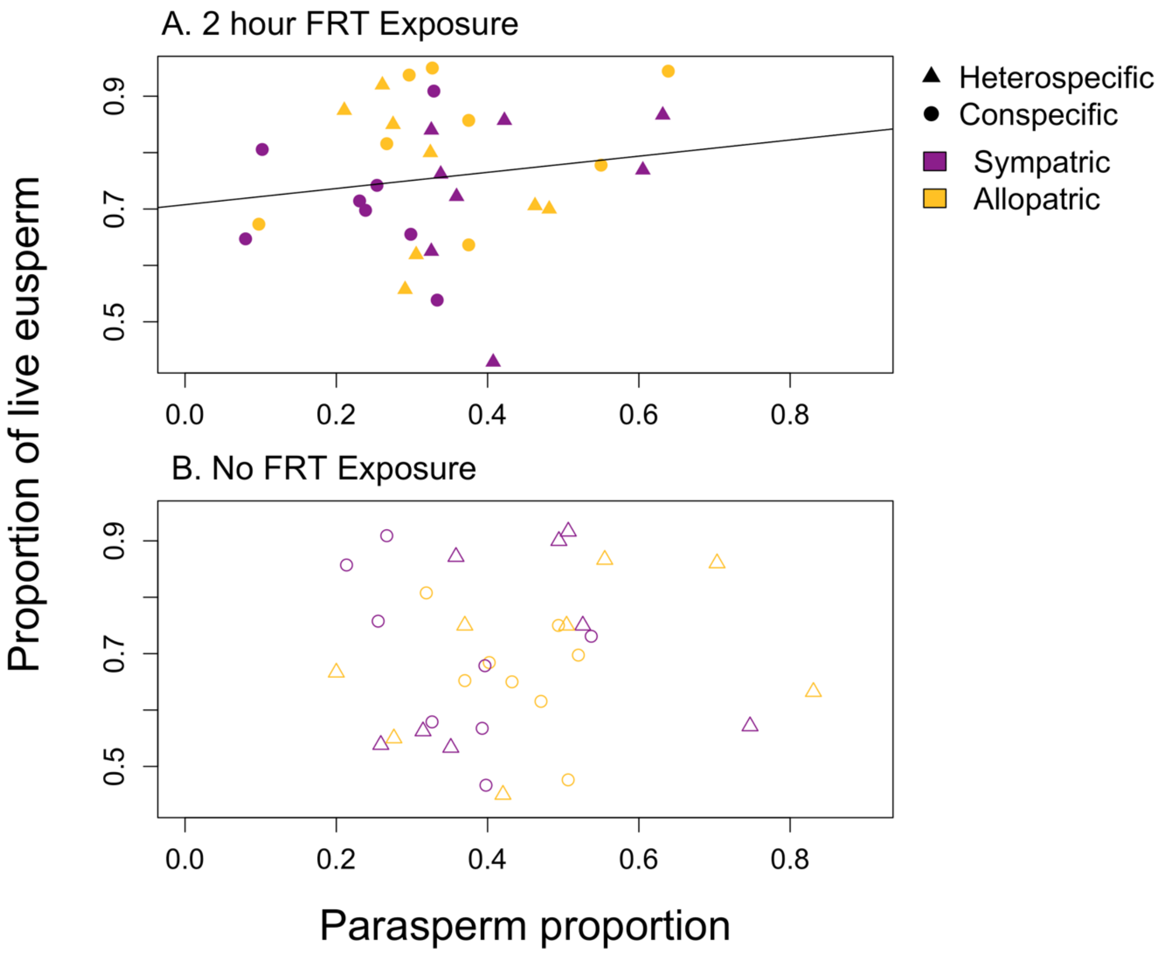


**Figure S6.** (A) Parasperm proportion predicts eusperm viability after two hours of FRT exposure (GLMM, $\beta = 1.50$, $P = 0.044$). (B) Parasperm proportion does not predict eusperm viability in control crosses, where sperm is dissected immediately after mating. This figure shows the raw data, as opposed to the model fitted values shown in Figure 3. Triangles represent heterospecific matings, circles represent conspecific matings.
